# T_RM_ Integrins CD103 and CD49a Differentially Support Adherence and Motility After Resolution of Influenza Virus Infection

**DOI:** 10.1101/2020.02.14.947986

**Authors:** Emma C Reilly, Kris Lambert Emo, Patrick M Buckley, Nicholas S Reilly, Francisco A Chaves, Hongmei Yang, Patrick W Oakes, David J Topham

## Abstract

Tissue resident memory CD8 T (T_RM_) cells are a unique immune memory subset that develops and remains in peripheral tissues at the site of infection, providing future host resistance upon re-exposure to that pathogen. In the pulmonary system, T_RM_ are identified through S1P antagonist CD69 and expression of integrins CD103/β7 and CD49a/CD29(β1). Contrary to the established role of CD69 on CD8 T cells, the functions of CD103 and CD49a on this population are not well defined. This study examines the expression patterns and functions of CD103 and CD49a with a specific focus on their impact on T cell motility during influenza virus infection. We show that the T_RM_ cell surface phenotype develops by two-weeks post-infection and that each integrin contributes a distinct function regulating CD8 T cell motility both *in vitro* and *in vivo*, with CD49a facilitating migration and CD103 limiting motility through tethering. These results demonstrate for the first time how CD103 and CD49a differentially impact adherence and migration in the tissue, likely affecting overall retention, maintenance of T_RM_, and host protection.

**Significance Statement:** Current influenza vaccination strategies require annual immunizations, with fairly low efficacy rates. One technique to improve protection against a greater breadth of influenza viruses is to elicit broadly cross-reactive cell-mediated immunity and generate a local population of cytotoxic T cells to respond to conserved regions of circulating viruses. However, this approach requires improved understanding of how these cells migrate within and attach to the tissue, in order to persist and offer long-term immunity. This study investigates how receptors on the T cell surface impact the cell’s ability to interact with the tissue and provide evidence for which of these receptors are essential for protection. Furthermore, these studies reveal functional *in vivo* mechanisms of cellular markers used to characterize T_RM_.

## Introduction

Tissue resident memory CD8 T cells (T_RM_) are a critical and unique component of the adaptive immune system (1-3). Unlike circulating memory cell populations (T_CM_ and T_EM_), they reside within peripheral tissues after clearance of a pathogen or immunological insult, and dictate downstream innate and adaptive responses upon reactivation (2, 4). In the absence of T_RM_ cells, hosts display increased susceptibility and illness severity when exposed to related pathogens (2). Current vaccination strategies, especially for influenza, focus on generating protective antibodies, and these approaches do not elicit effective CD8 T cell-mediated protection. Neutralizing antibodies classically bind external epitopes of pathogens, and on the influenza virion, target the head region of the rapidly evolving hemagglutinin proteins (5) (6) (7). In contrast, T_RM_ cells primarily respond to internally derived pathogen epitopes which are less prone to external pressures and mutation (8). The capability to induce localized memory T cells through vaccination will facilitate increased cross-protection against a greater breadth of influenza strains. However, we need to better understand many aspects of T_RM_ biology in order to optimize their generation and maintenance.

Parabiosis experiments established that the T_RM_ are a non-circulating subset (9, 10), but specific surface markers have been identified to minimize the need for this procedure. T_RM_ populations display phenotypic variation between tissue types, however, the majority express CD69 and integrins CD49a/CD29 and CD103/β7 to some degree (11, 12). Early studies suggested that CD69 was essential for the T_RM_ population in the skin and lung (13, 14); however, its contribution to T_RM_ maintenance was recently shown to function primarily in the kidneys, with less of a requirement in other peripheral sites (10, 15). In the lung, CD49a is necessary for the survival of T_RM_ cells, in part through engagement of its ligand collagen IV, which limits apoptosis (2, 16). Absence of CD49a results in increased susceptibility to a lethal challenge with a heterosubtypic influenza A virus, resulting from a more rapid decline in T_RM_ cell numbers. In other sites, lack of CD49a impacts the accumulation of gut-resident intraepithelial CD8 T cells (17) and blocking CD49a augments T cell positioning and the subsequent immune response (18). Eliminating the E-cadherin binding CD103 also results in a diminished population of T_RM_ cells in some tissues(13, 19-21). However, the CD103 requirement is less clear, and its functions *in vivo* may support initial accumulation of the cells rather than long-term persistence (21, 22).

In other cell types, integrins facilitate attachment, interactions with other cell populations, and support motility (23) (24). Early studies provide some insight into the contributions of CD49a and CD103 to the preservation of the T_RM_ population through blocking or deletion of the integrin (2) (19) (20) (21). These experiments establish a role for these integrins as markers essential for optimal T_RM_ maintenance, but do not delve into the direct interactions that lead to these functional ramifications. Therefore, we investigated how these integrins regulate interactions with the tissue through adherence and motility with the ultimate aim of determining whether these integrins equally contribute to host protection.

## RESULTS

### Pulmonary CD8 T cells Express CD49a and CD103 Early After Influenza Virus Clearance

T_RM_ cells are classically identified through expression of CD69 and integrins CD103 and CD49a. CD69 limits egress of T_RM_ cells from tissues by acting as an antagonist to S1P, and reduces responsiveness to S1P1 gradients that direct the cells towards the draining lymphatics (25, 26). The functions of the surface integrins CD103 and CD49a on T cells are less well appreciated. Additionally, the temporal development of this surface phenotype is not well characterized. Bona fide T_RM_ cells can be found in the airways, lung tissue, and trachea at 3 months post-infection (2). These cells express CD49a and/or CD103. We asked whether this phenotype arose early after virus clearance by comparing the integrin phenotypes at two weeks post-infection, with T_RM_ cells present at 3 months. CD8 T cells were examined in two different models of influenza virus infection. In the first, C57BL/6 mice were infected intranasally with H3N2 influenza A virus HKx31. In the second model, which is used for all imaging-based experiments described in this study, GFP OT-I cells were adoptively transferred prior to infection with an HKx31 variant expressing the OvaI SIINFEKL peptide in the neuraminidase stalk (27). To clearly separate cells in the tissue from cells circulating or intimately associated with the vasculature, antibody was introduced intravascularly prior to harvesting cells from the airways, tracheal tissue, or lung tissue (28, 29). Close to 100% of virus-specific CD8 T cells in the airways were negative for intravascular labeling, indicating that almost all of the cells are within the tissue parenchyma (Supplementary Figures 1 and 2). At 3 months post HKx31 infection, greater than 80% of NP/PA tetramer positive T cells in the airways express CD49a, with approximately 15% co-expressing both integrins (Figure 1A). This phenotype was comparable in the transfer system where airway OT-I cells were examined (Figure 1A). Only 10-50% of CD8 T cells in the lung were IV protected, depending on the time point evaluated (Supplementary Figure 1). For this reason, both the tissue and vasculature associated populations were examined within this organ. Approximately 20-30% of CD8 T cells within the tissue express only CD49a, with 5-30% expressing both integrins (Figures 1B). A subset of vascular CD8 T cells express CD49a alone, and only a small subset co-expresses CD103 (Figure 1B). Of note, within the CD49a positive subsets, tissue CD8 T cells have significantly higher levels of CD49a staining (Supplementary Figure 3.) Unexpectedly, cells in the trachea were almost exclusively IV protected and virus-specific CD8 T cells cells at 3 months post-infection were predominantly CD49a/CD103 double positive in both systems (Supplementary Figure 1 and Figure 1C).

**Figure 1.**
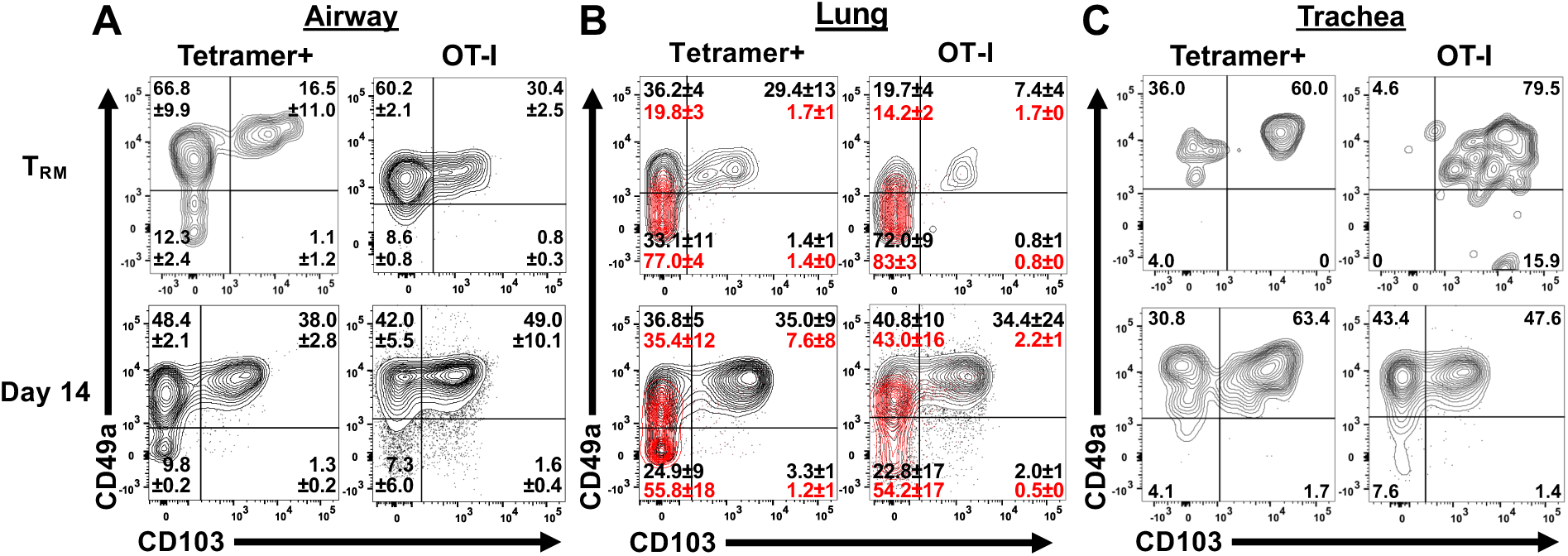
T_RM_ integrin phenotype arises by 2-weeks post-infection. Mice were infected with HKx31 influenza virus or OT-I CD8 T cells were adoptively transferred prior to infection with HKx31-OvaI (OT-I). Tissue virus-specific cells (black) or vasculature associated cells (red) were assessed by a combination of NP and PA tetramer (Tetramer+) or by gating on OT-I CD8 T cells (OT-I) at 3 months or 14 days post-infection. Cells were examined in the airways through bronchoalveolar lavage (A), Lung tissue (B), and Trachea (C). Results shown are mean +/- SD from 1 representative of 3 independent experiments.

Cells examined in wild type mice at day 14 post-infection demonstrated over 80% positivity for CD49a with a frequency of over 40% expressing CD103 and this phenotype was consistent with that in the OT-I transfer system (Figure 1A). Trachea and lung OT-I T cells within the tissue looked remarkably similar at this time point, with over 70% expressing CD49a and roughly one half of those cells also expressing CD103 (Figures 1B,C). Similar to the T_RM_ time point, a subset of vascular labeled CD8 T cells in the lungs expressed CD49a alone, however again the level of expression was significantly reduced compared with cells from the tissue proper. The high proportions and comparable integrin expression signatures at these time points suggest they are generated early after viral clearance, consistent with the postulation that T_RM_ develop during resolution and repair of the damaged airways.

### CD49a and CD103 Display Distinct Motility Functions *in vitro*

CD49a and CD103 have been shown to bind and interact with collagen and E-cadherin, respectively (30-32) (33, 34). However, in T cells, the understanding of how these interactions regulate motility is not fully established. Using a 2D *in vitro* system, we first asked the question, do CD49a and CD103 facilitate migration? On day 14 post-infection, virus-specific CD8 T cells identified through expression of GFP were imaged on mouse collagen IV or E-cadherin. GFP OT-I T cells on collagen IV displayed robust motility, with median speeds of 4.48 µm/min (IQR 3.85) displacement rates of 0.79 µm/min (IQR 2.29), and directionality as measured by straightness of 0.20 (IQR 0.33) (Figures 2A-D). Addition of CD49a blocking antibody resulted in reduced speeds, displacement, and straightness, suggesting that the migration was mediated through the CD49a-collagen IV interaction (Figures 2A-E and Supplementary Video 1). To verify that the observed effect was not indicative of deterioration in health of the cells, an antibody against CD103 was added to cells migrating on collagen. Blocking CD103 on collagen had no effect on motility, establishing that cells were migrating on collagen IV in a CD49a-dependent manner (Figure 2A-C,F). Cells plated on E-cadherin showed minimal displacement compared with cells on collagen IV (Figures 2A-C,H). The majority of cells remained fixed at a single point and addition of a CD103 blocking antibody did not show any overall changes in speed, displacement, or straightness, suggesting that CD103 does not support motility in this context (Figures 2A-C,H,I and Supplementary Video 2). Of note, blocking CD103 slightly decreased the number of adhered cells, likely due to detachment of some cells from the surface. Blocking CD49a on E-cadherin did not have any effect, compared with control cells (Figures 2A-C,H,J). To more closely examine these integrins engaging their respective substrates, cells were imaged on collagen IV and E-cadherin using TIRF microscopy. Cells on the collagen IV surface appeared elongated and even in the presence of tail-like structures, and continued to migrate (Figure 2G and Supplementary Video 3). In stark contrast, cells on E-cadherin often appeared flattened and had trailing tethers or tails, with far less displacement than is observed on the collagen substrate (Figure 2K and Supplementary Video 4). These data led us to the hypothesis that CD49a binding of collagen IV promotes migration and CD103 interacting with E-cadherin provides a point or points of attachment to limit overall motility.

**Figure 2.**
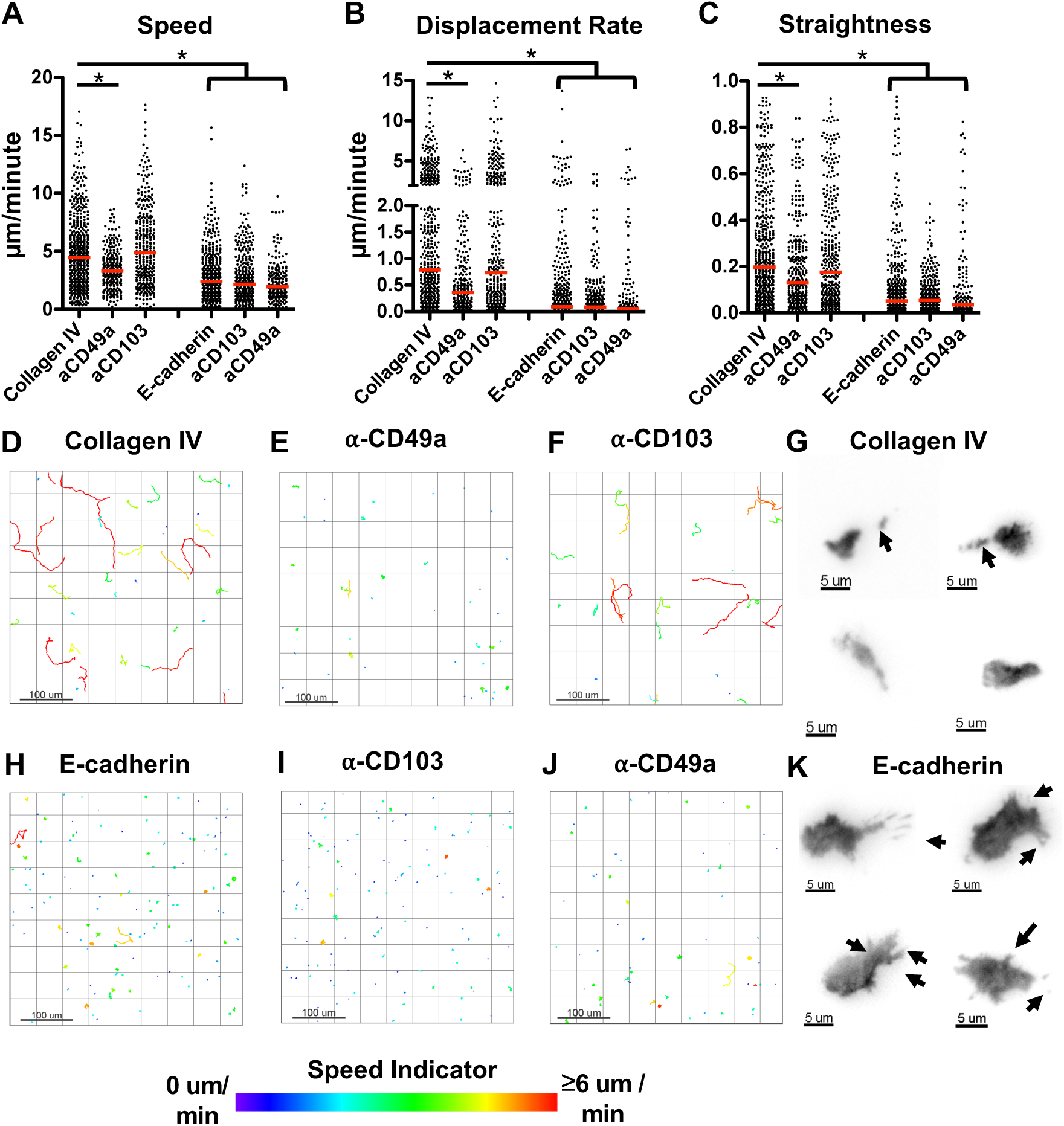
Day 14 CD8 T cells from lung migrate on collagen and tether to E-cadherin. OT-I T cells were extracted and negatively enriched from influenza infected lung at day 14 p.i. Cell motility was quantified on mouse collagen IV or E-cadherin before and after addition of blocking antibody. Speed (track length/time) (A), Displacement Rate (displacement/time) (B), and Straightness (total track length/displacement) (C) were calculated. Representative tracks from one experiment of 3-6 are shown with color indicating average track speed (D-F, H-J). Representative TIRF images on collagen and E-cadherin (G,K) are also shown. Data in A-C are from 6 control mice and 3 for each experimental condition. Red bar indicates the median. Significance was first established within each independent experiment and also with compiled data. *p<0.05

### CD49a and CD103 Are Unevenly Distributed on the Cell Surface and In Proximity to Respective Substrates in Tissue

We have shown that CD8 T cells can express CD49a and/or CD103 by flow cytometry and that these integrins provide distinct motility functions when examined *in vitro*. However, in both sets of experiments, cells were extracted from the tissues, which may not be representative of the entire population and does not indicate cellular location of the integrin (35). To identify the cellular localization of CD49a and CD103, tracheal tissue whole mounts from mice at days 14, 21, 42, and 3 months post-infection were stained and examined by microscopy. By day 14 post-infection CD49a could be identified in proximity to collagen as measured by second harmonic generation, directly underlying the epithelium (Figure 3A). Within the epithelial layer, predominantly CD103 expressing cells were identified (Figure 3A). At days 21 and 42 post-infection, it was clear that higher expression of these integrins on the cell surface was localized to specific foci and it was more common to see expression of a single integrin type at a given region (Figure 3B,C). This suggested that some cells may be utilizing only one interaction or that a cell could be interacting with multiple substrates at distinct cellular regions simultaneously. By 3 months post-infection, we could identify dual expression of both integrins at specific locations. Alternatively, at this time point we also observed CD103 expression on cellular protrusions embedded within the epithelium, supporting the hypothesis that CD103 binding of E-cadherin expressed in epithelial cell junctions can act as a tether to epithelial cells to limit movement (Figure 3D). We were similarly able to identify CD8^+^ cells in human trachea obtained from the LungMAP consortium, the Biorepository (1U01HL122700) and the LungMAP Data Coordinating Center (1U01HL122638). We observed CD103 expressing cells closer to the tip of the airway epithelial cells and dual expressing CD8^+^ cells closer to the basement membrane (Figure 3E), suggesting that the conclusions drawn from our system may at least partly translate to humans.

**Figure 3.**
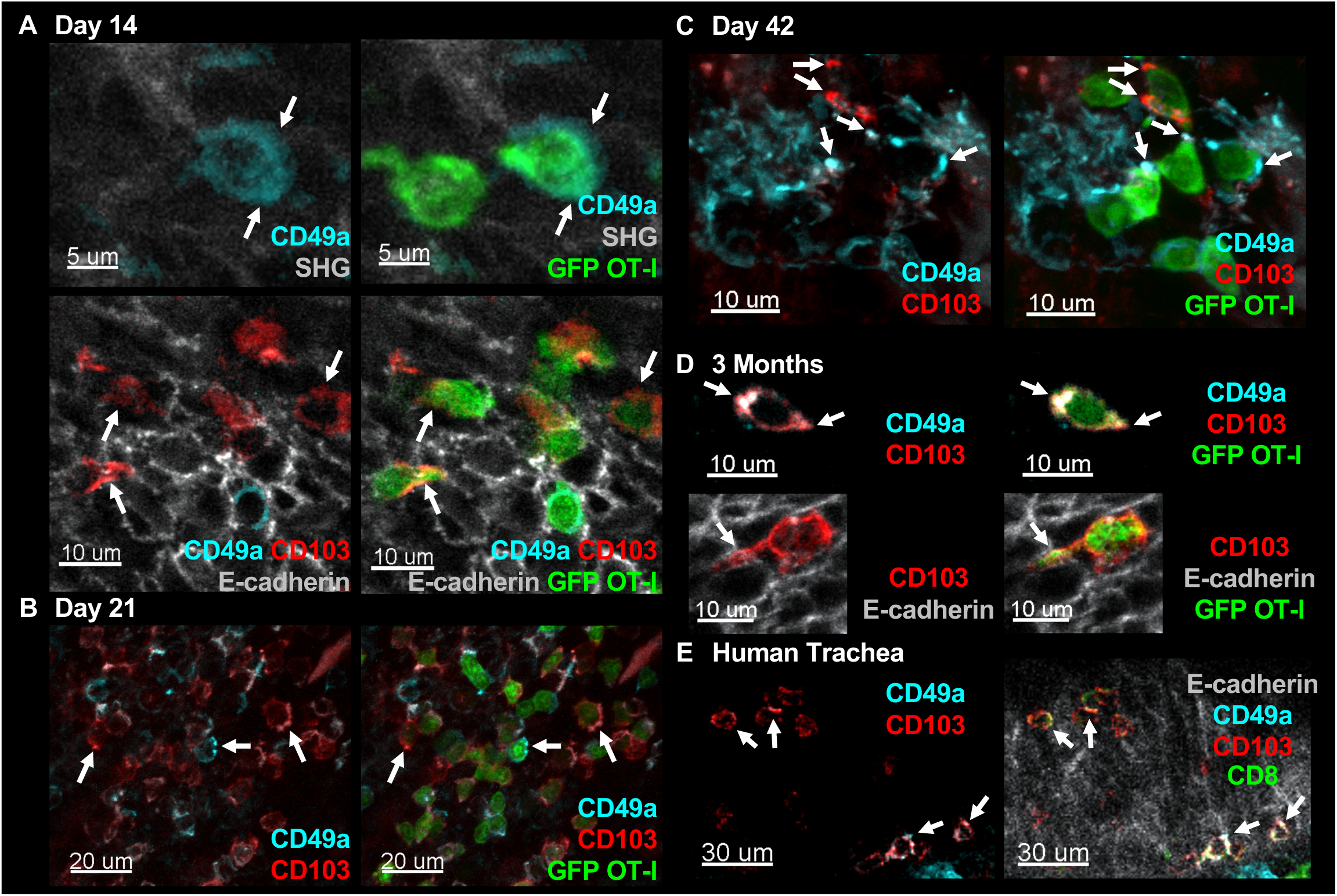
CD49a and CD103 are expressed more highly at specific regions of cells and can be seen in proximity to integrin substrates. Tracheal whole mounts from mice that received GFP OT-I CD8 T cells prior to infection were stained for CD49a and CD103 where indicated. Tracheas were examined at days 14 (A), 21 (B), 42 (C), and 3 months (D) post-infection. Human tracheal whole mount was stained with anti-human CD8, CD49a, and CD103 (E). Arrows indicate regions of localized staining.

### CD49a Promotes Motility and CD103 Limits Speed *in vivo*

One main goal of this study was to evaluate the role(s) of these integrins in cell motility *in vivo*. To achieve this, we utilized an intravital tracheal imaging system, previously developed in the lab (36). Transgenic OT-I T cells from WT or integrin deficient mice were transferred to wild type hosts and imaged at 2 weeks post-infection to examine CD8 T cell movement over time. In comparison to the simplistic 2D collagen system, wild type cells observed in the trachea displayed fairly low speeds (median of 1.95 µm/min with IQR of 1.82), without extensive displacement (median 0.33 µm/min with IQR of 0.41) (Figure 4A-D and Supplementary Video 5). This is likely due to the combination of signals the cells are receiving through multiple interactions and a consequence of being in a more confined environment. However, eliminating CD49a in GFP OTI cells resulted in further limited motility, consistent with the concept that CD49a is facilitating locomotion on collagen IV (Figures 4A-C,E and Supplementary Video 6). Conversely, absence of CD103 resulted in increased speeds compared with wild type cells (Figures 4A-C,F and Supplementary Video 7). The number of cells at this time point however was reduced, so we employed a second approach to examine the contribution of CD103 in this system. Wild type OT-I cells were transferred into naïve wild type hosts and starting at day 7 post-infection, mice were given a CD103 blocking antibody or isotype antibody every other day until imaging. To ensure that the antibody was reaching the cells, separate mice were examined for the presence of the antibody bound to virus-specific CD8 T cells, using an anti-rat IgG antibody. Cells in the trachea, lung, and airways of anti-CD103 treated mice were labeled with the anti-rat IgG, while Isotype control mice showed no binding of the secondary antibody (Supplementary Figure 4A). Absence of staining with the CD103 antibody prior to flow cytometric analysis suggested that not only was the in vivo antibody already bound, but that it was likely saturating. Blocking CD103 using this approach resulted in numbers of cells comparable to wild type at day 14 post-infection. However, the motility was markedly similar to that observed using the knockout cells. OT-I CD8 T cells displayed increased speeds in anti-CD103 treated mice compared with controls, reinforcing a role for CD103 in limiting motility (Figure 4G-J and Supplementary Video 8).

**Figure 4.**
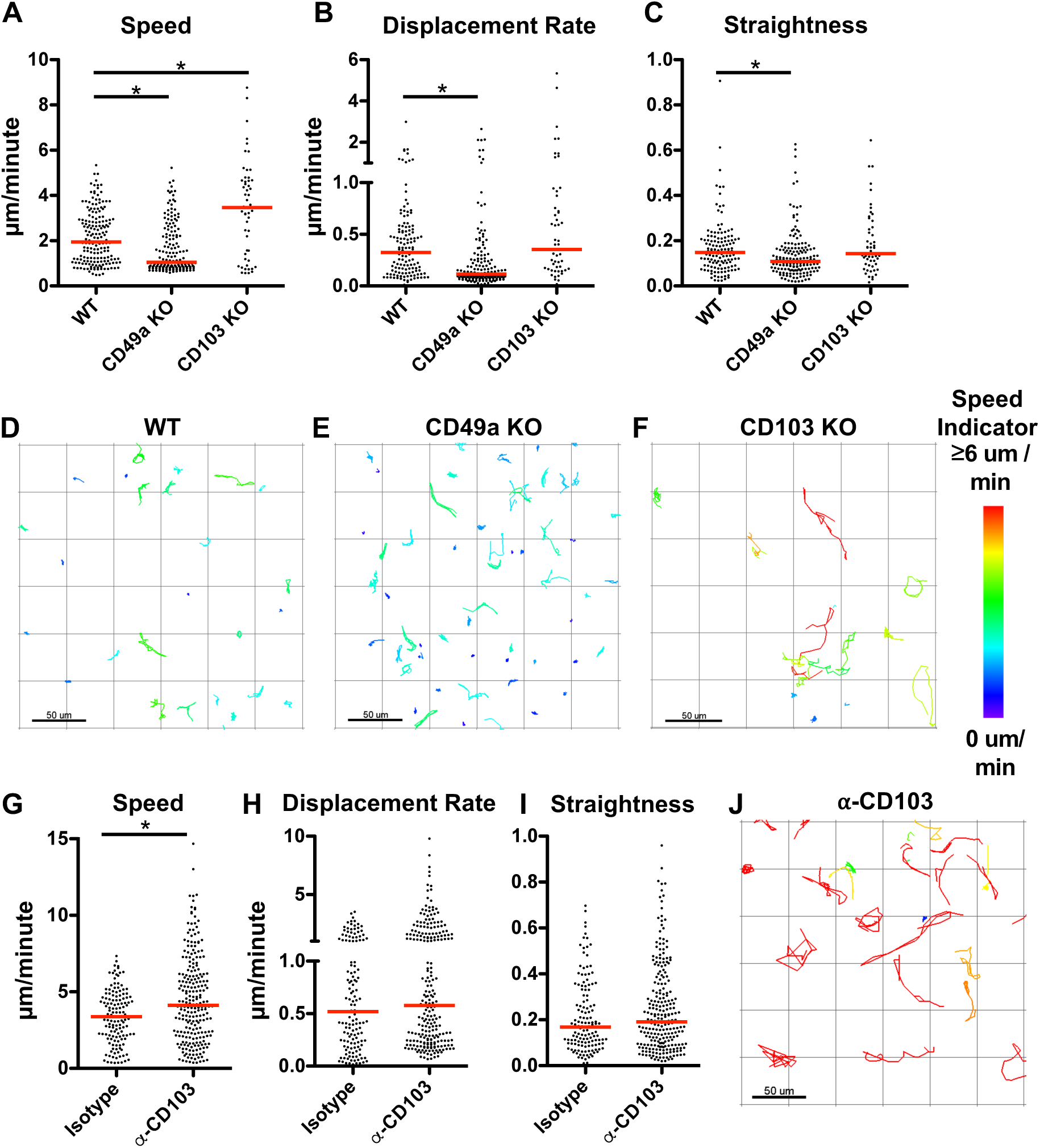
CD49a facilitates motility and CD103 limits speed. Integrin sufficient and deficient GFP OT-I CD8 T cells were adoptively transferred prior to infection with HKx31-OvaI influenza virus. Intravital imaging of the trachea was utilized to examine CD8 T cell motility *in vivo* on day 14 p.i. Speed (A), Displacement Rate (B), and Straightness (C) were enumerated. Representative tracks are shown with the color indicating the average track speed. Speed (G), Displacement Rate (H), Straightness (I) for CD103 blocking or isotype control treatment, and representative tracks with CD103 blocking antibody (J). Data were combined from 3-4 mice for each condition. *p<0.05

### CD49a and CD103 in T_RM_ Formation and Function

Both integrins have been implicated in formation of T_RM_ in the tissue. To determine whether the absence of one or the other integrin affects the overall integrin phenotype early after viral clearance, cells were examined using flow cytometry after transfer of either WT or integrin deficient cells at day 14 post-infection. In the absence of CD49a, no changes in the frequency or number of CD103 positive cells were observed (Figure 5A-B). Likewise, cells deficient in CD103 expressed similar frequencies of CD49a at this same timepoint (Figure 5C-D). The total number of CD103 deficient CD8 T cells expressing CD49a was slightly, but not significantly decreased (Figure 5D). However, this was not unexpected based on the *in vivo* data. Overall, our data suggest that the level of each integrin is regulated independently of the other. However, there could still be functional compensatory mechanisms.

**Figure 5.**
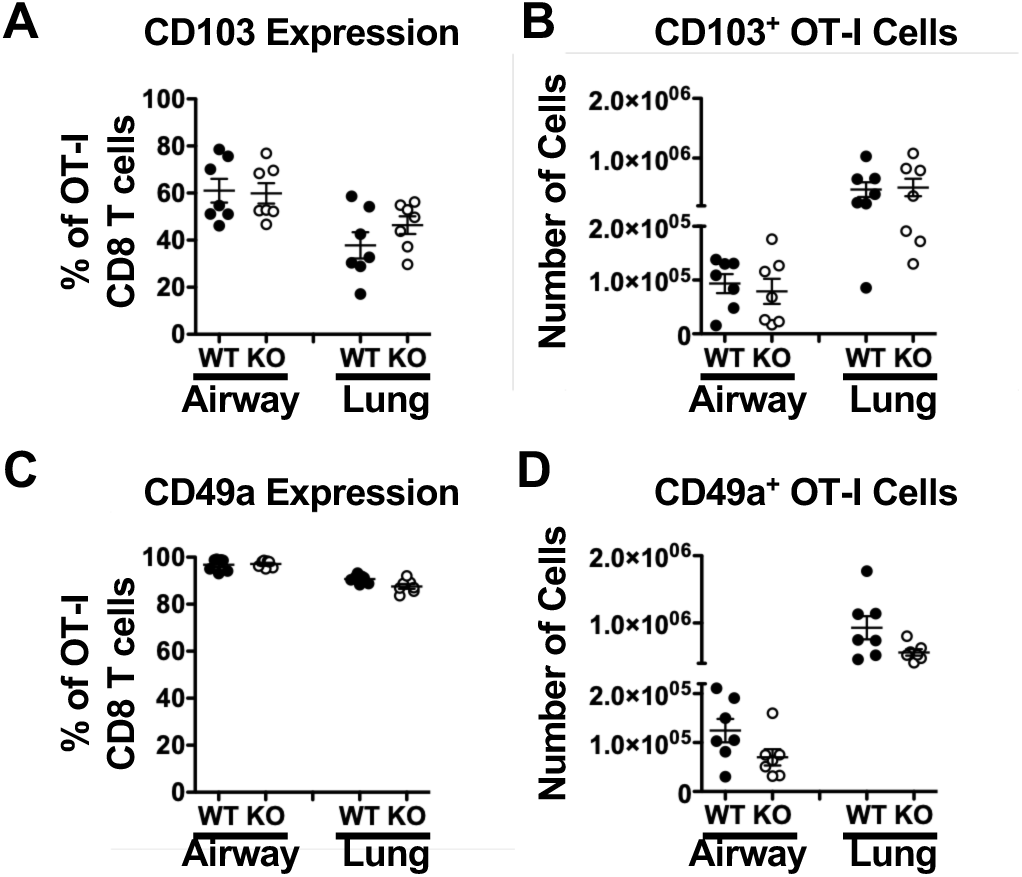
CD49a and CD103 expression is not influenced by the absence of the other integrin. WT or integrin deficient cells were transferred into WT mice prior to infection with HKx31-OvaI influenza virus. Airway (BAL) and lung were examined on day 14 p.i. for the presence of CD103 on WT and CD49a KO OT-I cells (A-B) or CD49a on WT and CD103 KO OT-I cells (C-D). Data are shown as percentage of the OT-I T cell population or absolute number of cells. Data shown are 2 experiments combined.

Therefore, we next sought to determine whether CD49a and CD103 could equally contribute to host defense. Our lab previously showed a requirement for CD49a to protect a host from heterosubtypic influenza virus challenge at 3 months post-infection (2). However, the contribution of CD103 in secondary protection has not been established. To examine this, WT and CD49a deficient mice were infected with an H3N2 virus HKx31 to generate a pool of memory cells. Three days and one day prior to reinfection with H1N1 virus PR8, both genotypes of mice were given either CD103 blocking antibody or PBS control. Again, to ensure localization of the antibody, separate mice were examined for binding of a secondary antibody (Supplementary Figure 4B). CD49a deficient mice, regardless of treatment, succumbed to the challenge by day 8, only one day later than non-immune mice with no prior infection (Figures 6A-B). WT mice given CD103 blocking antibody were protected equally as well as mice given control treatment, and the majority of mice survived the challenge (Figures 6A-B). These data show a clear requirement for CD49a, but further suggest that the CD103:E-cadherin interaction is not necessary at late time points for T_RM_-mediated host protection.

**Figure 6.**
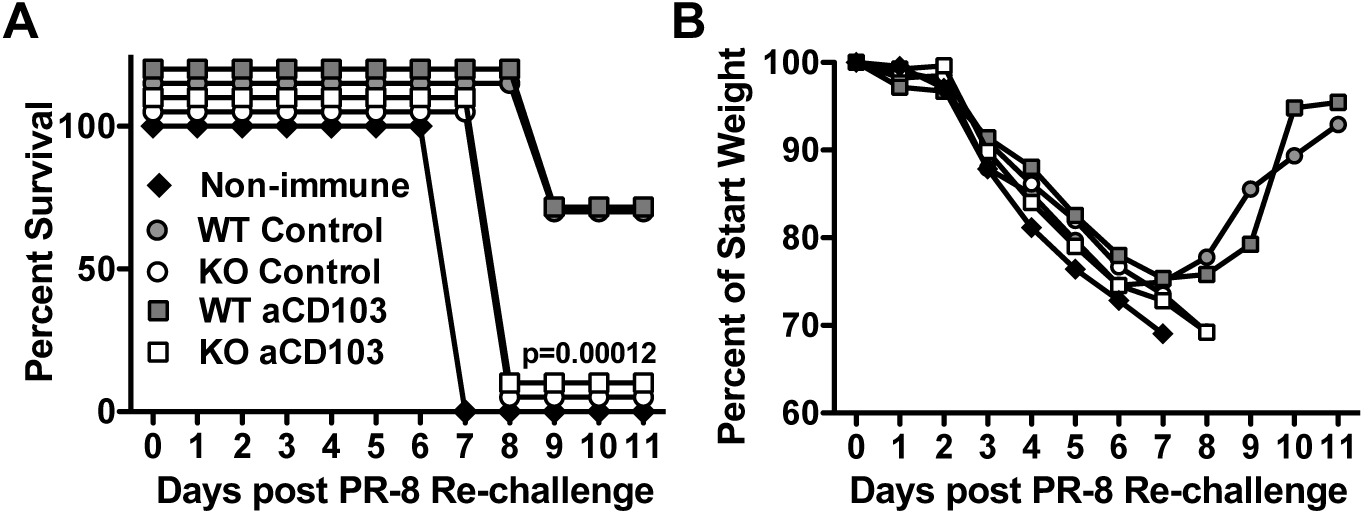
CD49a is critical for heterosubtypic protection. WT and CD49a KO mice were infected with HKx31 influenza virus. At 3 months post-infection, mice were given CD103 blocking antibody or PBS control. Mice were then infected with a lethal dose (1000EID_50_) of PR8 influenza virus. Non-immune mice were naïve at the time of PR8 infection. Mice were monitored daily for survival (A) and weight loss (B). Data shown are representative from 1 of 2 independent experiments. n=3-4mice/group and is representative of 2 independent experiments.

## Discussion

Integrins CD49a and CD103 appear on the surface of CD8 T cells in the airways, trachea, and lung tissue early after clearance of influenza virus, by day 14. While these integrins are viewed as markers of T_RM_ cells, they actually offer CD8 T cells the capacity to interact with collagen IV in the lamina densa underlying the epithelium, collagen I in the interstitium, and the epithelial cells directly through E-cadherin binding. The expression of CD49a and CD103 early after viral clearance suggest they are playing a critical role contributing to the development and maintenance of T_RM_. Previous work has established that active TGF-β, which is present during disease resolution, is essential for expression of CD103 and CD49a (37). In the gut, this has been highlighted by eliminating TGF-β signaling to T cells, which results in continued recruitment of effector cells, but no demonstrable population of tissue memory (37). Additionally, Nur77 signaling has recently been linked to direct regulation of CD49a transcription in endothelial cells potentially implicating a role for *in situ* TCR stimulation in modifying surface expression in this system (38, 39). Integrins can also be regulated through outside-in signaling, suggesting that direct interactions with collagen or E-cadherin on cells poised within or in proximity to the airway, may enhance surface expression of CD49a or CD103, respectively (40).

Distinct subsets of CD8 T cells in the airway, lung tissue, and trachea can be identified through expression of CD49a and/or CD103. Early characterization of T_RM_ cells focused exclusively on CD103 and CD69 as identifiers of the population, however, our lab has shown that CD49a is critical for long-term maintenance of the population in the lungs, in part through survival signals received via interactions with collagen IV (2) (16). CD103 appears to regulate early recruitment and persistence, but is less critical after resolution of infection for maintenance of the memory pool (21) (22). We sought to examine the outcomes of these integrins interacting with their ligands both *in vitro* and *in vivo* to reveal potential mechanisms regulating the T_RM_ population and ultimately host protection.

*In vitro* imaging of virus-specific CD8 T cells on collagen IV and E-cadherin suggested that CD49a and CD103 function differently in motility and adherence. CD8 T cells migrated readily on collagen IV in a CD49a-dependent manner. While previous reports examining other cell types such as fibroblasts and endothelial cells suggested that CD49a’s function was primarily to adhere to collagen, the differences observed could be attributed to distinct mechanisms mediating adhesion between cell types (38, 41, 42). Inhibition or knockdown of CD49a in tumor cells resulted in decreased migration compared with their CD49a sufficient counterparts (43, 44). Similarly, human CD8 T cells *ex vivo* were unable to migrate through a collagen IV coated transwell when CD49a was blocked (45). More extensive examination of whether these CD8 T cells on collagen require active re-engagement of ligand to survive long term is warranted with the knowledge that the CD49a:collagen IV interactions are important for cell persistence. Additionally, it would be of interest to determine whether these outcomes are specific to collagen IV, or if these phenomena can be more broadly applied to other collagens, such as collagen I. Alternatively, T cells on E-cadherin displayed a strong attachment, with limited migration observed. This fundamentally different role in regulation of cell motility is consistent with both lymphoid and non-lymphoid cells examined in other organs such as the gut(33, 34), and may mediate local retention of cells at the peripheral site and provide continued surveillance.

Within the tissue, CD49a and CD103 are not uniformly distributed on the cell surface. In fact, the expression of each integrin was often observed in regions of the integrin ligand. CD103 coincided with E-cadherin expression within the epithelium, consistent with the capacity to interact with epithelial cells. CD49a could be identified in close proximity to collagen directly underlying epithelial cells, suggesting it may use this surface to explore other regions of the epithelium and receive pro-survival signals. To determine whether these integrins played similar roles *in vivo* to what was observed *in vitro*, we used a combination of genetically deficient T cells and blocking antibodies to examine the motility of virus-specific CD8 T cells in the trachea of 2-week infected mice. Consistent with cell motility observed in the 2D setting, absence of CD49a *in vivo* resulted in further arrest of the cells, indicating a role in cell locomotion. This observation conflicted with our initial interpretation that CD49a was important for retention by physically adhering the cells to the matrix. Instead, these studies reveal that CD49a is critical for cell motility, a feature that is not previously associated with retention. In contrast, knocking out or blocking CD103 resulted in increased speeds of the cells examined, suggesting a reduced ability to tether to epithelial cells. While *in vivo* migration of T cells lacking CD49a has not been extensively examined, transplanted human skin in a mouse model of psoriasis showed that CD49a blockade limited the ability of T cells to position within the epidermal layer and sequestered these cells to the dermis, signifying an important role for CD49a in motility and/or positioning (45). Diametrically, imaging of CD103 deficient CD8 T cells in the skin during a mouse model of herpes simplex virus infection displayed increased speeds, supporting a role for CD103 in tethering to the skin epithelium and limiting overall motility (20).

For years, the CD8 T cell field has relied on using integrins CD103 and CD49a to identify the T_RM_ subset; however, our understanding of the functions of these receptors in regulating interactions with the tissue and T cell motility are incomplete and underappreciated. In this report we examined the roles of CD103 and CD49a in the CD8 T cell subset present after clearance of viral infection. Absence or blocking of CD49a in this system showed an unexpectedly strong effect, indicating that other integrins such as CD51 which confer motility in CD4 T cells, are not sufficient to compensate in this population (46). Alternatively, our data suggest that CD103’s main function is not to mediate locomotion, but rather is critical for interactions with epithelial cells, effectively tethering them to the epithelial surface and facilitating maintenance and surveillance. γδ T cells in the gut have been shown to “floss” the epithelium (47), and we propose T_RM_ in the airways exhibit a similar behavior. Of note, while the effects observed were consistent both *in vitro* and *in vivo*, our analysis examined the complete pool of virus-specific CD8 T cells present in the tissue. Future experiments with more focused imaging and transcriptomics analyses will be needed to dissect how different integrin phenotypes regulate interactions within the tissues and their resulting cellular functions. It will also be of interest to determine how these integrins affect motility in response to secondary infection, and further extend these findings to other organ systems and infections. Given the ubiquity of the ligands for CD49a and CD103 at mucosal and epithelial barriers, our data support that the mechanisms described here may extend beyond the pulmonary system and be critical for secondary immune protection at boundary surfaces throughout the body.

## Materials and Methods

### Mice

All mice were maintained and managed in university approved, pathogen free facilities using microisolator technology. C57BL/6 mice were purchased from Jackson Laboratories (Bar Harbor, ME). All mice were 8–10 weeks at the start of the infection. A colony of OT-I transgenic mice that express a TCR specific for the OVA SIINFEKL (OVA_257-264_) peptide presented in the context of H-2K^b^ were crossed with a transgenic mouse expressing GFP under a β-actin promoter maintained in house at the University of Rochester. This line of mice was crossed with VLA-1 KO mice (2) or CD103 KO mice (B6.129S2(C)-Itgae^tm1Cmp^/J) (Jackson Laboratories). This study was carried out in strict accordance with the recommendations in the Guide for the Care and Use of Laboratory Animals as defined by the National Institutes of Health. Animal protocols were reviewed and approved by the Institutional Animal Care and Use Committee (IACUC) of the University of Rochester. All animals were housed in a centralized and AAALAC-accredited research animal facility that is fully staffed with trained husbandry, technical, and veterinary personnel.

### Treatment and cell transfer

For imaging experiments, 1×10^6^ splenocytes from WT or integrin KO GFP OT-I cells were transferred intravenously on day −1 to naïve C57BL/6 hosts. WT and integrin KO mice were checked by flow cytometry to ensure that comparable numbers of CD8 T cells were being transferred. For CD103 blocking, mice were given 150μg anti-CD103 antibody (clone M290) or isotype control antibody (clone 2A3) IP in a 200μL volume. Mice were given injections on days 7, 9, 11, and 13. For blocking prior to secondary challenge, mice were given 150μg anti-CD103 or PBS on days −3 and −1 and 150μg IN on day −2. For cellular preparations, 0.2μg CD8β-FITC, CD45-PE, or CD45-Brilliant Violet 786 was given intravenously in 100uL PBS 3 minutes prior to organ harvest.

### Viruses and Infections

The influenza H3N2 A/Hong Kong/X31 (HKx31) virus, HKx31-OVAI influenza virus that expresses the ovalbumin (OVA257-264 SIINFEKL) peptide in the neuraminidase viral protein, and H1N1 A/Puerto Rico/8 (PR8) were grown and titered in embryonated chicken eggs and harvested as allantoic fluid preparations (27). Mice were sedated with avertin and infected IN with 10^5^ EID_50_ of HKx31, 3×10^3^ EID_50_ HKx31-OVAI, or 10^3^ EID_50_ PR8 in 30μL of PBS. After infections in all experiments including survival studies, mice were monitored daily for weight loss, ability to ambulate, ability to intake food and water, and signs of discomfort including ruffled fur, hunched posture, and guarding behavior.

### Cellular Preparations

Bronchoalveolar lavage (BAL) samples were obtained by cannulating the trachea and flushing lungs with 1x PBS. Cells were spun down and lysed with 500μL of ACK lysis buffer (Ammonium-Chloride-Potassium) for 5 minutes at room temperature. Cells were washed in PBS containing 1% FBS (PBS serum) and resuspended for counting in 1mL of PBS serum. Lungs and trachea were perfused with 1x PBS, removed, and separated into trachea and right and left lobes. Lung and trachea tissue were dissociated in C tubes by the GentleMACS (Miltenyi Biotek) using the Lung01.02 program. Samples were incubated in 5mL [2μg/mL] Collagenase II (Worthington) in RPMI +8% FBS at 37°C for 30 minutes, with gentle agitation. After digestion, samples were further dissociated using the Heart01.01 program. Cell suspensions were filtered through 100μm filters prior to 75:40 Percoll (GE Healthcare) discontinuous gradient separation. The top layer, containing fat and other debris, was removed by aspiration. The cell layer was harvested and washed, prior to counting and staining. Counting was achieved through Trypan blue exclusion on a hemocytometer.

### Flow cytometry

Single cell suspensions were stained in PBS serum, purified CD16/32 (clone 2.4G2), and some combination of the following antibodies: TCRβ, CD8α, CD69, CD49a, CD103, CD44, CD62L. All antibodies were obtained from either Biolegend or BD Biosciences. NP and PA tetramers were obtained from the NIH tetramer core facility (Atlanta, GA). Cells were also stained with fixable live/dead aqua indicator in PBS (Invitrogen). Cells were analyzed by LSRII (BD Biosciences) in the University of Rochester Flow Cytometry core facility and analyzed using FlowJo software (Tree Star).

### In vitro imaging

Cell migration chambers (Millicell EZ slide eight-well glass; Millipore; or Delta T dish; Biotech) were prepared by coating their glass bottom with [10 µg/cm^2^] mouse collagen IV (Corning) in 0.05M HCl for 1 hour at room temperature or [2.5 µg/cm^2^] mouse E-cadherin (R&D Systems) overnight at 4°C in DPBS. Prior to imaging, plate was washed 3x in PBS. For *in vitro* migration imaging, GFP OT-I cells were negatively enriched from lungs for CD8 T cells using the Miltenyi Biotek CD8 Isolation Kit (#130-104-075) and resuspended in L15 medium (Invitrogen) containing [2mg/mL] D-glucose. These cells were placed in the chamber at 37°C, allowed to adhere for 20 minutes, and microscopy was conducted using a TE2000-U microscope (Nikon) coupled to a CoolSNAP HQ CCD camera with a 20× objective (CFI Plan Fluor ELWD DM; Nikon) and 0.45 numerical aperture. Cells were imaged for 20 minutes with images taken every 20 seconds. Blocking antibodies (1µg) were added directly into the media and cells were subsequently imaged for 20 minutes. All images were acquired in NIS Elements (Nikon). Migration analysis was performed in Imaris software (Bitplane). For TIRF microscopy, images were collected using a 60x 1.49 NA oil immersion TIRF objective (Nikon) place on an inverted Ti-E microscope (Nikon) coupled to a Dragonfly TIRF module (Andor). Multi-field time lapses were taken while samples were kept at 37C with 5% CO_2_ with a stage top incubation system (Oko-Lab).

### Tracheal Whole Mount Imaging

Excised mouse trachea was incubated in PBS serum, anti-CD16/32, and indicated antibodies for 4 hours at room temperature. The following antibodies were used in different combinations at the indicated dilutions: CD103-AlexaFluor 594 (Biolegend) 1:50, CD49a-AlexaFluor 647 (BDBiosciences) 1:25, E-cadherin-APC (Biolegend) 1:50. Images were acquired on an Olympus FV1000 (Center for Advanced Light Microscopy and Nanoscopy Shared Resource Laboratory) using an Olympus Plan Apo 60x/1.43NA objective. Fresh, transplant donor-quality human tracheal tissue was obtained from the Human Tissue Core of the Lung Development Molecular Atlas Program (LungMAP) at the University of Rochester. The University of Rochester IRB approved and oversees this study (RSRB00047606). Human trachea was stained for 72 hours in PBS serum and the following antibodies: hCD103-AlexaFluor 594 (Biolegend) 1:40, hCD8-AlexaFluor 488 1:40, hCD49a-AlexaFluor 647 (Biolegend) 1:40, E-cadherin-APC (Biolegend) 1:40. Human tissue was washed in PBS and fixed in 4% Paraformaldehyde for 2 hours prior to mounting. Mouse tissue was washed in PBS and mounted unfixed. All whole mount tissue sections were mounted using Fluoromount G (SouthernBiotech).

### Intravital Multiphoton Imaging

Intravital tracheal imaging was performed as described in Lambert Emo, *et al (36)*. Briefly, mice were anesthetized with [65mg/kg] pentobarbital. Hair was removed from one hind leg, exposing skin for MouseOX Plus sensor (Starr Life Sciences) monitoring. The hair was removed from the thoracic area with scissors and the mouse was placed on a heated stage. Once surgical plane of anesthesia was determined by both lack of pedal and palpebral reflex, the coat was opened between the chin and the top of the rib cage. The submandibular salivary glands were separated to reveal the muscles covering the trachea and these muscles were separated to expose the trachea. A small flexible plastic support was placed under the trachea to separate it from the surrounding muscle, tissue, and coat. A small incision was made between cartilage rings below the larynx. An 18 gauge steel cannula was inserted into the opening in the trachea until just below the sternum. Mice were ventilated through the cannula using the Harvard Inspira ASV ventilator with 100% O2 and 0.5% Isoflurane according to weight. The MouseOX Plus thigh sensor was attached to the exposed thigh and mouse was monitored throughout imaging session. Oxygenation levels were maintained at >95% and heart rate between 250 and 600 beats per minute. The rectal body temperature was monitored and maintained using a small animal feedback regulated heating pad. After animal was stable and verification of lack of both pedal and palpebral reflex, panucronium bromide [0.4mg/kg] was administered intramuscularly. Agarose (0.05%) was added to exposed trachea and plastic wrap was used to create a water basin for the objective.

All images were collected by an Olympus FVMPE-RS system (Olympus, Center Valley, PA) using Olympus 25× water objective (XLPLN25XWMP2, 1.05NA). The system was equipped with two two-photon lasers: Spectra-Physics InSightX3 (680nm-1300nm, Spectra-Physics, Santa Clara, CA) and Spectra-Physics MaiTai DeepSee Ti:Sapphire laser (690nm-1040nm). There were four Photon Multiplier Tubes (PMTs) and two filter cubes (Blue/Green cube: 420-460nm/495-540nm, red/fRed cube: 575-630nm/645-685nm) for multi-color imaging. Galvonometer scanner was used for scanning, and all images were acquired at ∼1frame/s. PMT gains for all imaging were used between 550 and 700 a.u. in the Olympus Fluoview software. All images were registered within Matlab (MathWorks) using the registration code provided by the Center for Integrated Research Computing (https://github.com/TophamLab/Registration) and were analyzed and visualized using Imaris (Bitplane) software.

### Statistics

The majority of statistics were performed using Prism Analysis Software (GraphPad). Flow cytometry data groups were compared using a Student’s T test and are shown as mean and SD. For imaging data which cannot be assumed to be normally distributed, the Kruskal Wallis Test was performed followed by Dunn’s comparisons with the control group. MFI data was compared with a paired T-test. Significance was considered a p-value <0.05. Imaging data values are presented as median and interquartile range (IQR). For survival data a non-parametric Kaplan-Meier method was applied.

## Acknowledgements

The authors would like to thank the Flow Cytometry Core, the Multiphoton Core, and the Confocal and Conventional Microscopy Resource at the University of Rochester. The authors also thank the Center for Integrated Research Computing (CIRC) at the University of Rochester for providing computational resources and technical support. The human data shown in this manuscript was made possible through the LungMAP Consortium [U01HL122642], the Biorepository (1U01HL122700), and the LungMAP Data Coordinating Center (1U01HL122638) that are funded by the National Heart, Lung, and Blood Institute (NHLBI). The authors would also like to thank the Kim laboratory at the University of Rochester for generously allowing us to utilize their microscope for *in vitro* imaging studies. This work was supported by NIH/NIAID grant P01-AI102851.

## Figure Legends

**Supplementary Figure 1.**
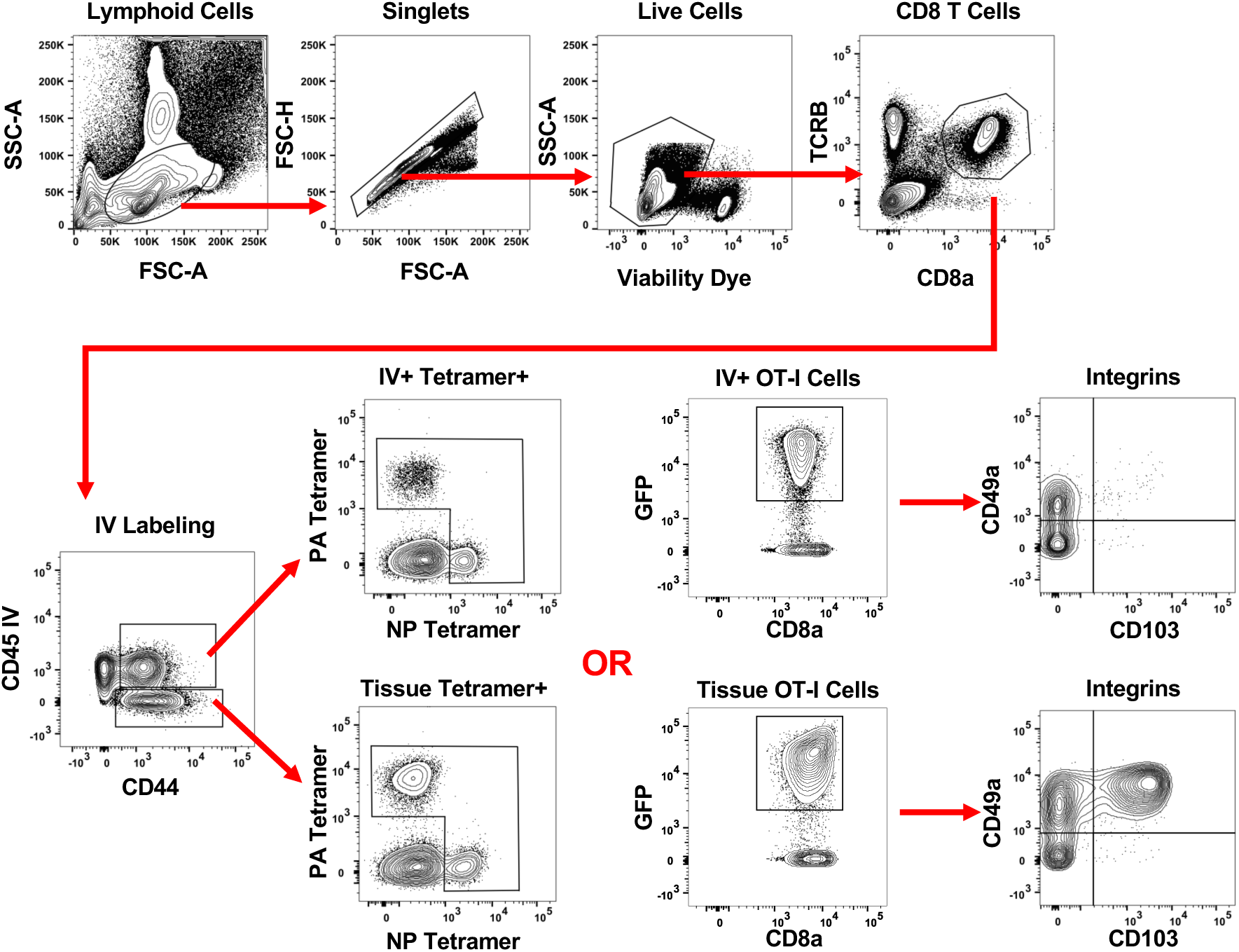
Gating strategy for identification of virus-specific CD8 T cells. Cells were first gated on lymphoid cells, singlets, and live cells. CD8 T cells were identified by TCRB and CD8a expression. These cells were examined for IV labeling and CD44 expression. The IV+ and IV-populations expressing CD44 were then examined for either tetramer staining OR GFP indicating OT-I T cells. These populations further examined for CD49a and CD103.

**Supplementary Figure 2.**
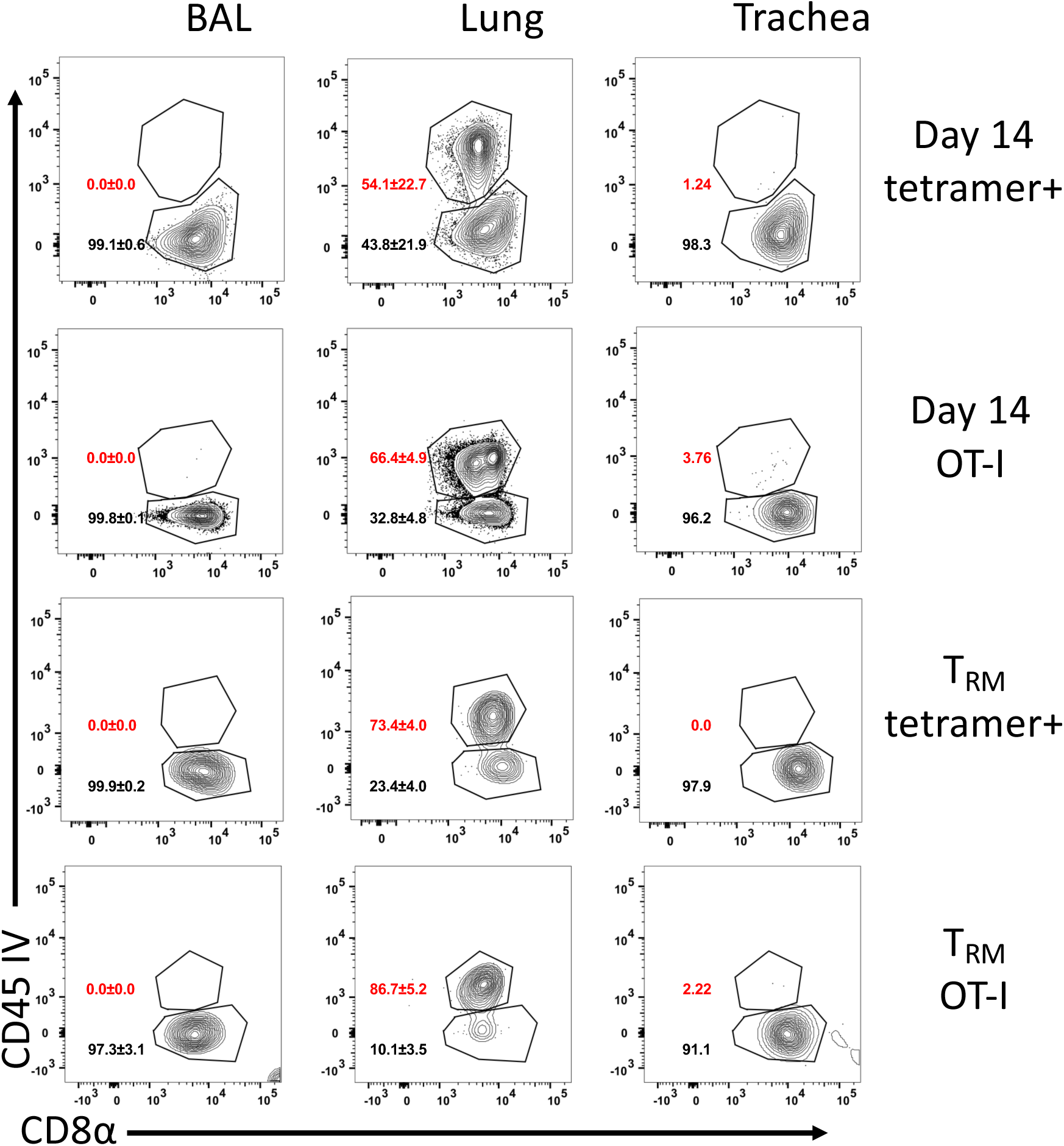
Intravascular labeling. IV labeling of virus-specific CD8 T cells was also examined in BAL, Lung, and Trachea at day 14 and 3 months (T_RM_) in the WT system and after transfer of OT-I T cells.

**Supplementary Figure 3.**
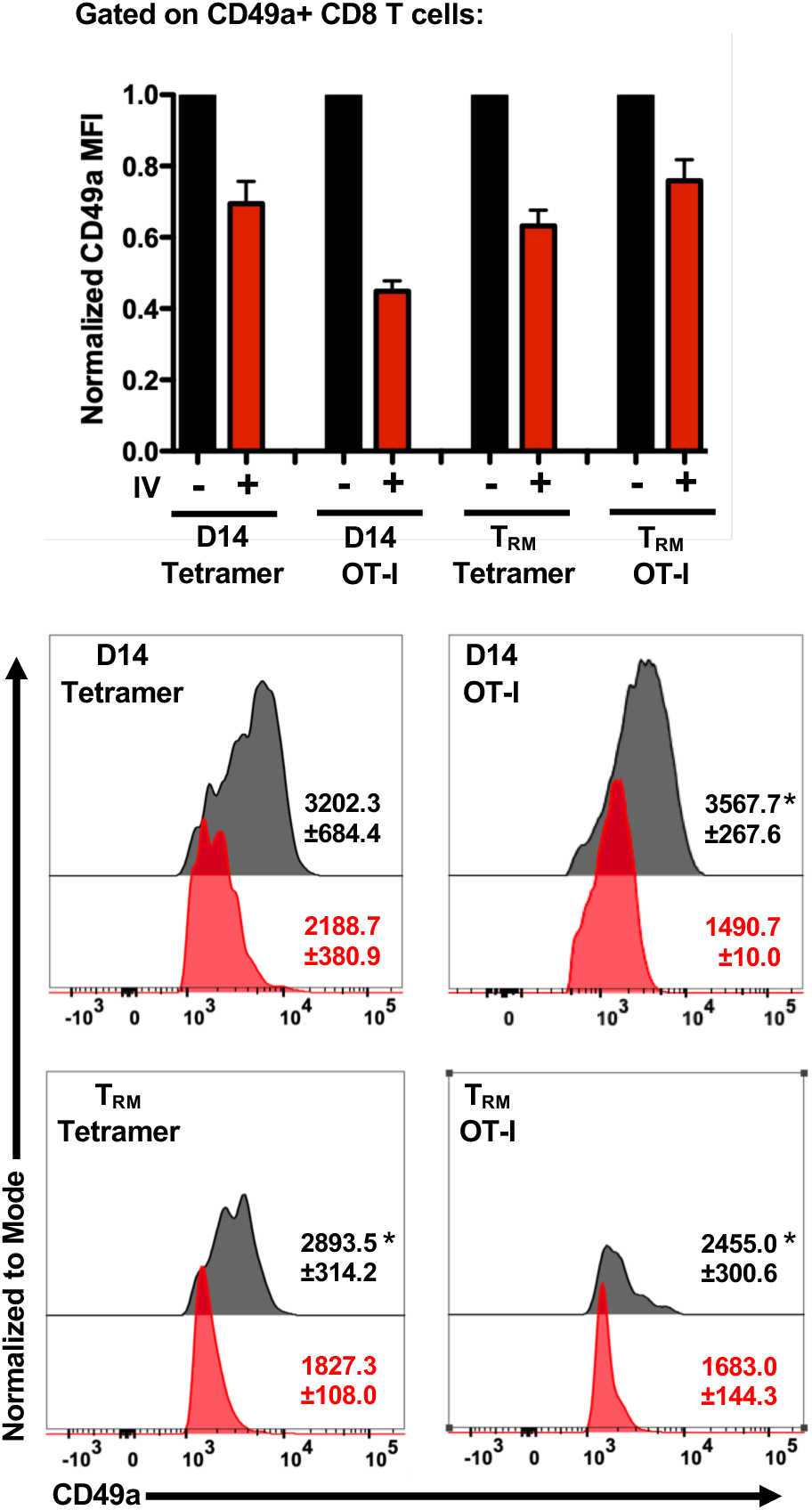
The Median Fluorescence Intensity of CD49a is significantly higher on cells within the tissue compared to vasculature. The CD49a+ subsets of tissue and vasculature associated virus-specific cells were examined for median fluorescence intensity of CD49a.

**Supplementary Figure 4.**
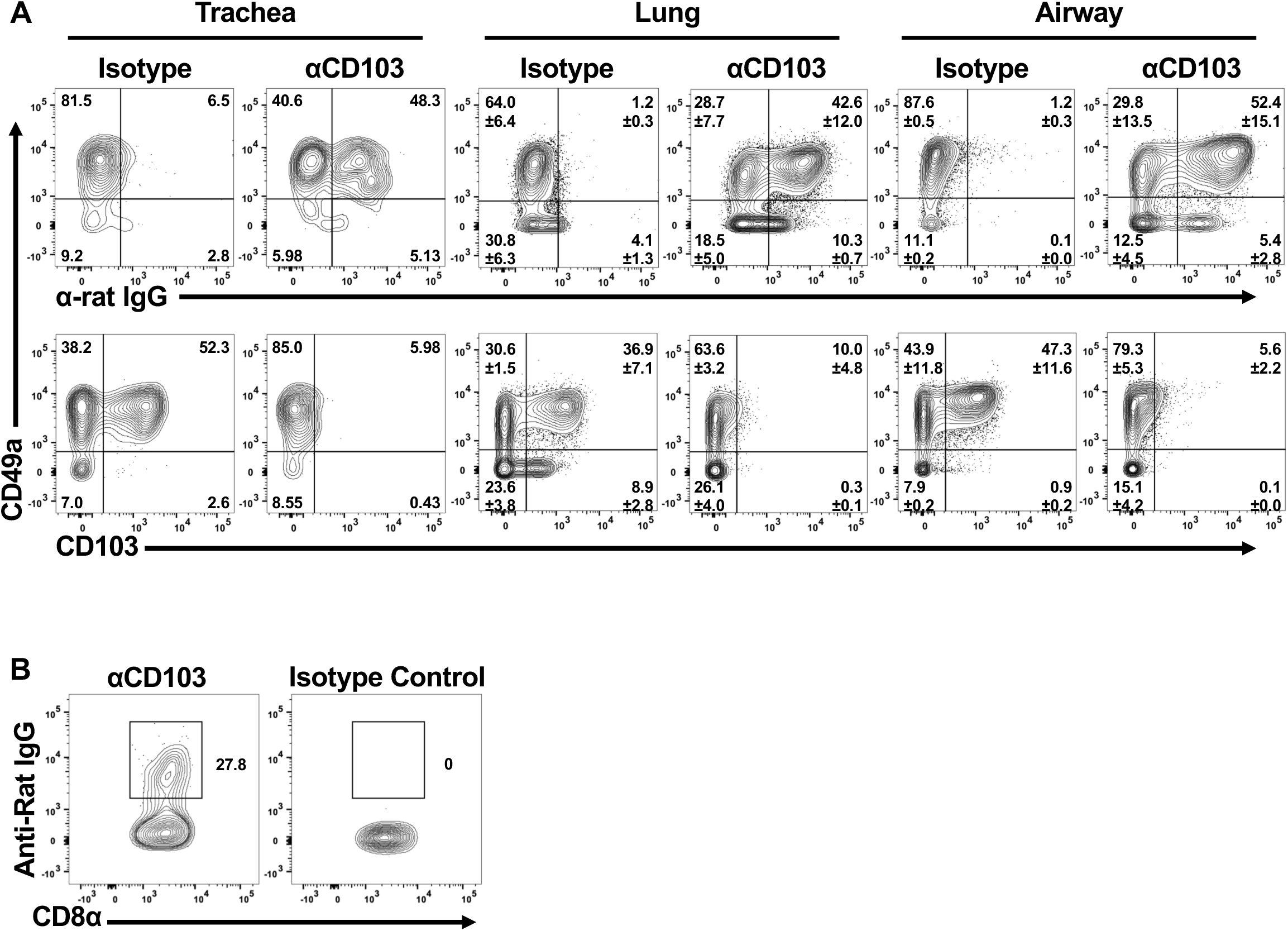
CD103 antibody given IP and or IN/IP is maintained on CD8 T cells. Anti-CD103 of Isotype control antibody was given to mice as described in materials and methods. Organs were harvested and cells were stained first with an anti-rat IgG antibody. After washing, cells were stained for other markers including CD103. Cells were examined at day 14 (A) or at memory time points prior to reinfection (B).

**Supplementary Video 1.** *In vitro* imaging of day 14 GFP OT-I T cells on collagen IV before and after administration of blocking CD49a antibody. The color of tracks are indicators of mean track speed (purple = 0um/min and red is > or = to 6um/min).

**Supplementary Video 2.** *In vitro* imaging of day 14 GFP OT-I T cells on E-cadherin before and after administration of blocking CD103 antibody. The color of tracks are indicators of mean track speed (purple = 0um/min and red is > or = to 6um/min).

**Supplementary Video 3.** TIRF imaging of a day 14 GFP OT-I T cells on collagen IV.

**Supplementary Video 4.** TIRF imaging of a day 14 GFP OT-I T cells on E-cadherin.

**Supplementary Video 5.** *In vivo* imaging of wild type GFP OT-I cells in the trachea at day 14 post-infection. The color of tracks are indicators of mean track speed (purple = 0um/min and red is > or = to 6um/min).

**Supplementary Video 6.** *In vivo* imaging of CD49a deficient GFP OT-I cells in the trachea at day 14 post-infection. The color of tracks are indicators of mean track speed (purple = 0um/min and red is > or = to 6um/min).

**Supplementary Video 7.** *In vivo* imaging of CD103 deficient GFP OT-I cells in the trachea at day 14 post-infection. The color of tracks are indicators of mean track speed (purple = 0um/min and red is > or = to 6um/min).

**Supplementary Video 8.** *In vivo* imaging of wild type GFP OT-I cells in the trachea at day 14 post-infection of a mouse given CD103 blocking antibody. The color of tracks are indicators of mean track speed (purple = 0um/min and red is > or = to 6um/min).

